# The HuMet Repository: Watching human metabolism at work

**DOI:** 10.1101/2023.08.08.550079

**Authors:** Patrick Weinisch, Johannes Raffler, Werner Römisch-Margl, Matthias Arnold, Robert P. Mohney, Manuela J. Rist, Cornelia Prehn, Thomas Skurk, Hans Hauner, Hannelore Daniel, Karsten Suhre, Gabi Kastenmüller

## Abstract

The human metabolism constantly responds to stimuli such as food intake, fasting, exercise, and stress, triggering adaptive biochemical processes across multiple metabolic pathways. To understand the role of these processes and disruptions thereof in health and disease, detailed documentation of healthy metabolic responses is needed but still scarce on a time-resolved metabolome-wide level.

Here, we present the HuMet Repository, a web-based resource for exploring dynamic metabolic responses to six physiological challenges (exercise, 36 h fasting, oral glucose and lipid loads, mixed meal, cold stress) in healthy subjects. For building this resource, we integrated existing and newly derived metabolomics data measured in blood, urine, and breath samples of 15 young healthy men at up to 56 time points during the six highly standardized challenge tests conducted over four days. The data comprise 1.1 million data points acquired on multiple platforms with temporal profiles of 2,656 metabolites from a broad range of biochemical pathways. By embedding the dataset into an interactive web application, we enable users to easily access, search, filter, analyze, and visualize the time-resolved metabolomic readouts and derived results. Users can put metabolites into their larger context by identifying metabolites with similar trajectories or by visualizing metabolites within holistic metabolic networks to pinpoint pathways of interest. In three showcases, we outline the value of the repository for gaining biological insights and generating hypotheses by analyzing the wash-out of dietary markers, the complementarity of metabolomics platforms in dynamic versus cross-sectional data, and similarities and differences in systemic metabolic responses across challenges.

With its comprehensive collection of time-resolved metabolomics data, the HuMet Repository, freely accessible at https://humet.org/, is a reference for normal, healthy responses to metabolic challenges in young males. It will enable researchers with and without computational expertise, to flexibly query the data for their own research into the dynamics of human metabolism.

## Introduction

The human body continually adapts and dynamically responds to physiological perturbations and challenges, such as dietary intake, physical activity, or stress ^1^. On a molecular level, this response translates to corresponding changes in biochemical processes that produce or consume low molecular weight organic compounds (metabolites) like glucose, cortisol, or cholesterol. Metabolomics techniques based on mass spectrometry (MS) or nuclear magnetic resonance spectroscopy (NMR) can measure the levels of hundreds of these metabolites simultaneously in accessible body fluids such as blood or urine. These metabolite profiles provide a snapshot of a person’s metabolic state at a given time ^2^. A series of such snapshots taken at multiple time points during or immediately after a specific challenge (e.g., extended fasting, exercise, or fat- or carbohydrate-rich meals) allows monitoring the systemic metabolic adaptation to the challenge in a time-resolved manner, i.e., metabolomics enables us to watch metabolism ‘at work’.

An individual’s body is metabolically flexible and reacts to challenges swiftly to restore metabolic homeostasis. Impaired metabolic flexibility is a hallmark of many metabolic disorders, such as type 2 diabetes (T2D) and cardiovascular diseases ^3^. It leads to aberrations from the ‘normal,’ healthy response to challenges in patients. For example, the efficacy of an individual’s insulin-regulated response is compromised in T2D, which leads to delayed clearance of excess glucose from the blood after carbohydrate ingestion. The oral glucose tolerance test (OGTT) is a standardized challenge test to recognize such abnormalities in glucose response for early diagnosis of T2D. This test focuses on the dynamics of a single blood metabolite – glucose – as a readout of response, and a single challenge – the ingestion of 75 g glucose – to trigger a metabolic response. In principle, this concept of testing resilience to a metabolic stressor can be expanded to include a broader spectrum of metabolites using metabolomics (e.g., additionally monitoring lipids during an OGTT ^4–6^), different challenges (e.g., monitoring glucose after different meals ^7^), and a combination of both, to ultimately recognize metabolic aberrations in diseases or even before the onset of clinical symptoms.

As a prerequisite for the practical, clinical use of a broader set of metabolites and challenges, atypical responses to these challenges must be identified and differentiated from the normal adaptation of metabolism in a healthy state. To this end, detailed knowledge of metabolism’s typical, healthy dynamics and its variance across individuals is crucial. However, most metabolomics studies involving challenges collect only two samples, one before and one after (or under) the challenge. Only few studies collected time-resolved metabolomic profiles more precisely describing the ‘normal’ dynamic responses to standardized and all-day metabolic stressors, including different nutritional challenges ^8–11^ or exercise ^11–13^. The HuMet study ^11^, was specifically designed to capture the normal dynamics of metabolism in a homogenous, healthy group (n=15) on a high temporal resolution and across multiple challenges, including four highly standardized (i.e., reproducible) nutritional challenges, a physical exercise, and a stress test. While sample collection covered many time points, biofluids, and challenges, the initially obtained time-resolved metabolomic profiles of the HuMet study participants comprised mainly metabolites from amino acid and various lipid classes. In contrast, the studies by Moreville et al. and Contrepois et al. investigated only responses on exercise in blood but used a non-targeted metabolomics approach, more broadly covering human metabolism ^12, 13^.

Following the open science paradigm, availability of data representing normal metabolic dynamics has increased in public repositories such as MetaboLights ^14^ and Metabolomics Workbench ^15^. Nonetheless, accessing and leveraging time-resolved metabolomics data remains challenging, particularly for researchers who are not used to handling big and complex longitudinal datasets. Various tools, including the MetaboAnalyst ^16^ and R packages such as MetaboLousie ^17^, mixOmics ^18^, and OmicsLonDA ^19^ can help to process and analyze time-resolved metabolomics data. However, these tools and repositories are primarily designed for bioinformatics experts. Thus, usage of available datasets still requires knowledge and additional effort on the computational side, which puts a large burden on researchers for answering single *ad hoc* question such as *“Do blood levels of a metabolite of interest (e.g., a potential biomarker) change postprandially or in response to exercise? If so, when are its levels back to baseline in healthy individuals?”* based on these data. To lower the burden of data access and usage for the broader scientific community, these complex time-resolved data must be embedded into intuitively browsable public resources.

We here describe the *HuMet Repository*, a public online resource that allows intuitive, interactive exploration and visualization of a comprehensive time-resolved metabolomics dataset capturing the normal dynamics of metabolism in men. For building this resource, we re-examined samples from the HuMet study ^11^ using five complementary non-targeted mass spectrometry-based metabolomics and lipidomics methods. As a result, the HuMet Repository contains temporal profiles for, in total, 2,656 metabolites (thereof >2,100 from the new analyses) measured in blood, urine, and breath samples from 15 healthy young males who engaged in the 4-day HuMet trial with six different metabolic challenges and samples collected at up to 56 time points for each participant. For each challenge, we identify metabolites and groups of metabolites that change using univariate statistics as well as data-derived metabolic networks. A dedicated web-based interface enables users to search, browse and visualize this complex dataset for further explorative analysis without specific computational expertise. In three showcases, we exemplify the use of the HuMet Repository: (i) Applying the implemented search functions, we identify metabolites with similar trajectories, e.g., metabolites that show a steady decrease over the study phase and presumably stem from exposures before the study. (ii) We check the similarity of trajectories of the same metabolites determined on two different metabolomics platforms, providing insights into the concordance of these measurements. (iii) Making use of data-derived and knowledge-based metabolic networks, we inspect the dynamic changes during extended fasting across the whole metabolism and we compare the responses observed in selected pathways between three different meal tests (glucose, mixed, high-fat).

In summary, the interactive HuMet Repository represents a unique resource of time-resolved metabolomics data in response to different physiological challenges within the same healthy homogenous population while covering a wide variety of metabolomics approaches. The HuMet Repository is freely accessible at (https://humet.org/).

## Results

For building the HuMet Repository as a comprehensive resource of time-resolved metabolomic profiles in healthy individuals (**Supplementary Table 1**), we first analyzed 840 blood and 240 urine samples of the HuMet study participants on the non-targeted Metabolon HD4 metabolomic platform (Metabolon Inc., Durham, USA) ^20^, measuring metabolites from a broad spectrum of metabolite classes (**Supplementary Table 2**). This added time-resolved measurements for more than 2,100 metabolites to the previously reported data (limited to mainly amino acids and lipids), resulting in a total of 1.1 million data points within the complete HuMet dataset.

Based on these data, we identified significant metabolic responses to four nutritional challenges (extended fasting, intake of standardized liquid meals with different macronutrient compositions (glucose (OGTT), mixed (SLD), lipid-rich (OLTT)), exercise and stress. To provide a metabolome-wide overview of triggered responses, we mapped all metabolite changes onto metabolic networks which we reconstructed from the time-resolved data.

We then embedded the complete data and results into a computational framework, providing a web-based graphical user interface for data access and exploration (**Figure 1**). The web-interface is split into four modules that are tailored to (i) select individual metabolites or groups thereof according to various criteria (Module *Selection*); (ii) visualize temporal metabolic trajectories (Module *Time course*); (iii) provide an overview of results from statistical tests (Module *Statistics*); and (iv) integrate metabolites into networks to obtain a holistic view on pathways altered in response to the challenges (Module *Networks*).

**Figure 1.**
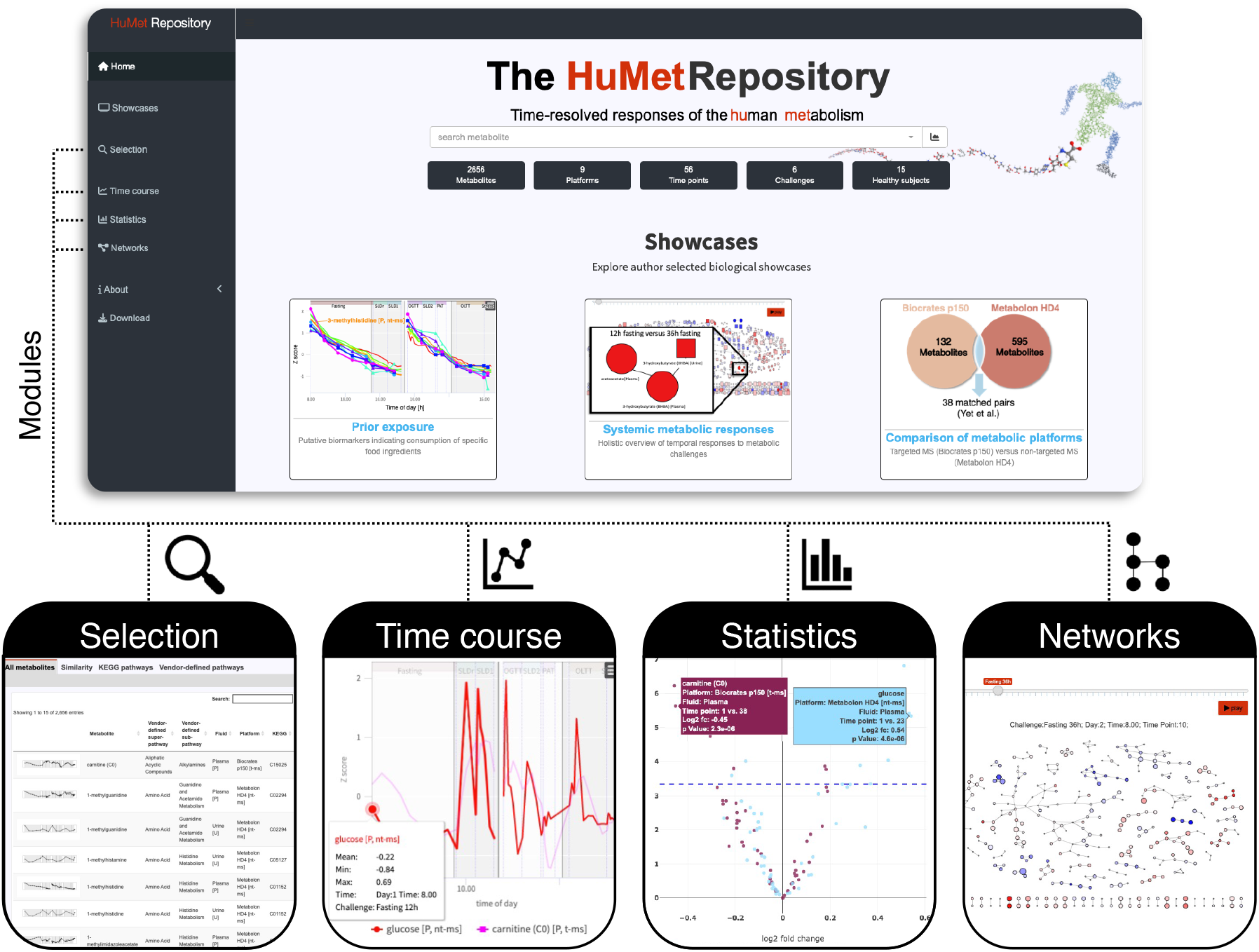
HuMet Respository frontend. The HuMet Repository (https://humet.org) integrates four individual modules to help explore the time-resolved metabolomics data of the HuMet study, reflecting responses to physiological challenges in healthy individuals. In the module *Selection*, the user can select metabolites from a table, with options for sorting and filtering by metabolite properties, including time course similarities. Line plots within the module *Time course* visualize time-resolved metabolite profiles of participants providing multiple options for data transformation and representation. Plots depicting statistical results from multiple analyses can be viewed in the module *Statistics*. The module *Networks* offers a holistic overview of metabolite changes within pre-defined and reconstructed biological pathways.

Finally, we present three showcases to exemplify applications of the HuMet Repository for exploratory analyses.

### Deep metabotyping provides time-resolved profiles for 2,656 metabolites

Using four non-targeted, mass spectrometry-based [nt-ms] analytical methods (Metabolon HD4 platform), we re-examined the 840 (15 subjects x 56 time points) plasma and 240 (15 subjects x 16 time points) urine samples of HuMet. This resulted in time-resolved relative quantifications for 595 and 619 metabolites in plasma and urine, respectively (**Table 1**). These metabolites span eight different metabolite classes called ‘super-pathways’ (amino acids, carbohydrates, cofactors and vitamins, energy, lipids, nucleotides, peptides, xenobiotics) and more than 83 different metabolic pathways (‘sub-pathways’). In addition, samples of four participants were analyzed on the Lipidyzer^TM^ platform, yielding quantifications of 965 molecules that provide structurally detailed information on complex lipids (see **Methods**).

**Table 1.**
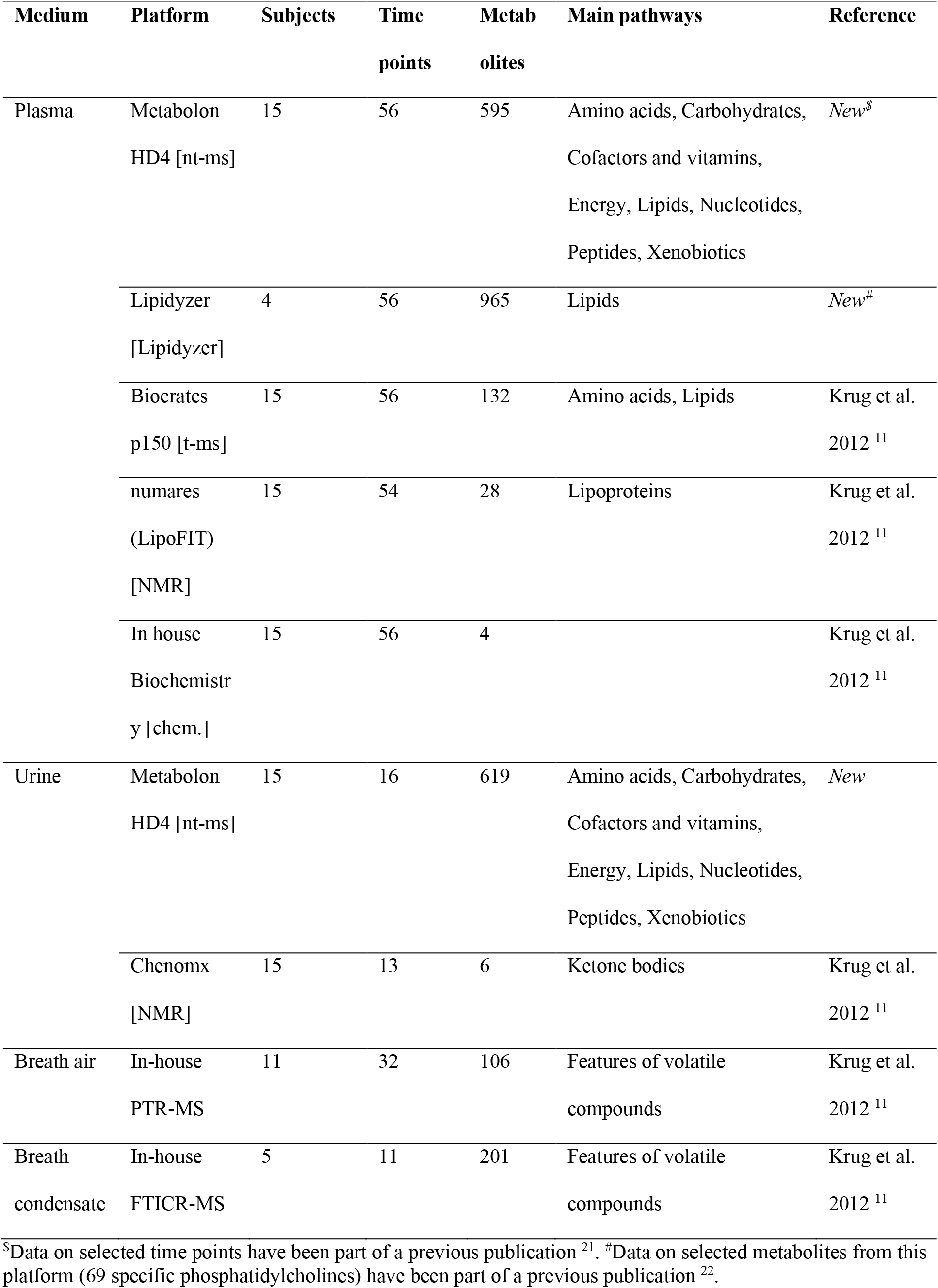
Overview of metabolomics data provided within the HuMet Repository.

In the HuMet Repository, we also included the previously published data from the initial metabolomics analysis of the HuMet samples ^11^, which covered mainly amino acids, lipids (acylcarnitines, glycerophospholipids, sphingolipids) and lipoproteins. The plasma concentrations of these metabolites were measured using the commercially available Biocrates p150 kit for targeted MS-based [t-ms] analysis. Levels of plasma lipoproteins were assessed at numares AG (formerly LipoFIT, Regensburg, Germany) applying an NMR-based approach. Moreover, breath air and breath condensate samples had been analyzed on in-house platforms of partners from academia. This first wave of measurements resulted in quantifications for 477 metabolomic measures for HuMet samples (**Table 1**). Descriptions of all (newly and previously measured) metabolites are provided in **Supplementary Table 2**. In total, the HuMet Repository, thus, provides access to time-resolved data for 2,656 metabolites in plasma, urine, and breath, covering a broad spectrum of metabolic pathways.

### Metabolic responses to six physiological challenges

To characterize the ‘normal’, healthy dynamics in metabolism under all-day physiological challenges, we analyzed the concentration changes of each metabolite during/after each of the six physiological challenges, to which the participants of the HuMet study were exposed during the four days of sample collection (**Figure 2**): In a first block of two days, participants had to fast for 36 h (*Fasting*) and were allowed to recover from fasting after breakfast and a lunch consisting of a standardized drink that represents a mixed meal (*SLDr, SLD1*). The second block of the study, which was conducted after a four-week break, included a physical activity test (*PAT*), a stress test (*Stress*), and three different nutritional challenges, namely an oral glucose tolerance test (*OGTT*) resembling a diet rich in carbohydrates, an oral lipid tolerance test (*OLTT*) resembling a high-fat diet, and the ingestion of the same liquid diet as used for the recovery from fasting (*SLD2*), resembling a mixed meal (for details see **Methods** and Krug et al. ^11^). Throughout the experiment, three different sample types (plasma, urine, and breath) were collected at up to 56 time points in variable time intervals (15 min - 2 h, depending on the challenge), enabling temporal profiling of metabolite changes during or after the six challenges for each participant.

**Figure 2.**
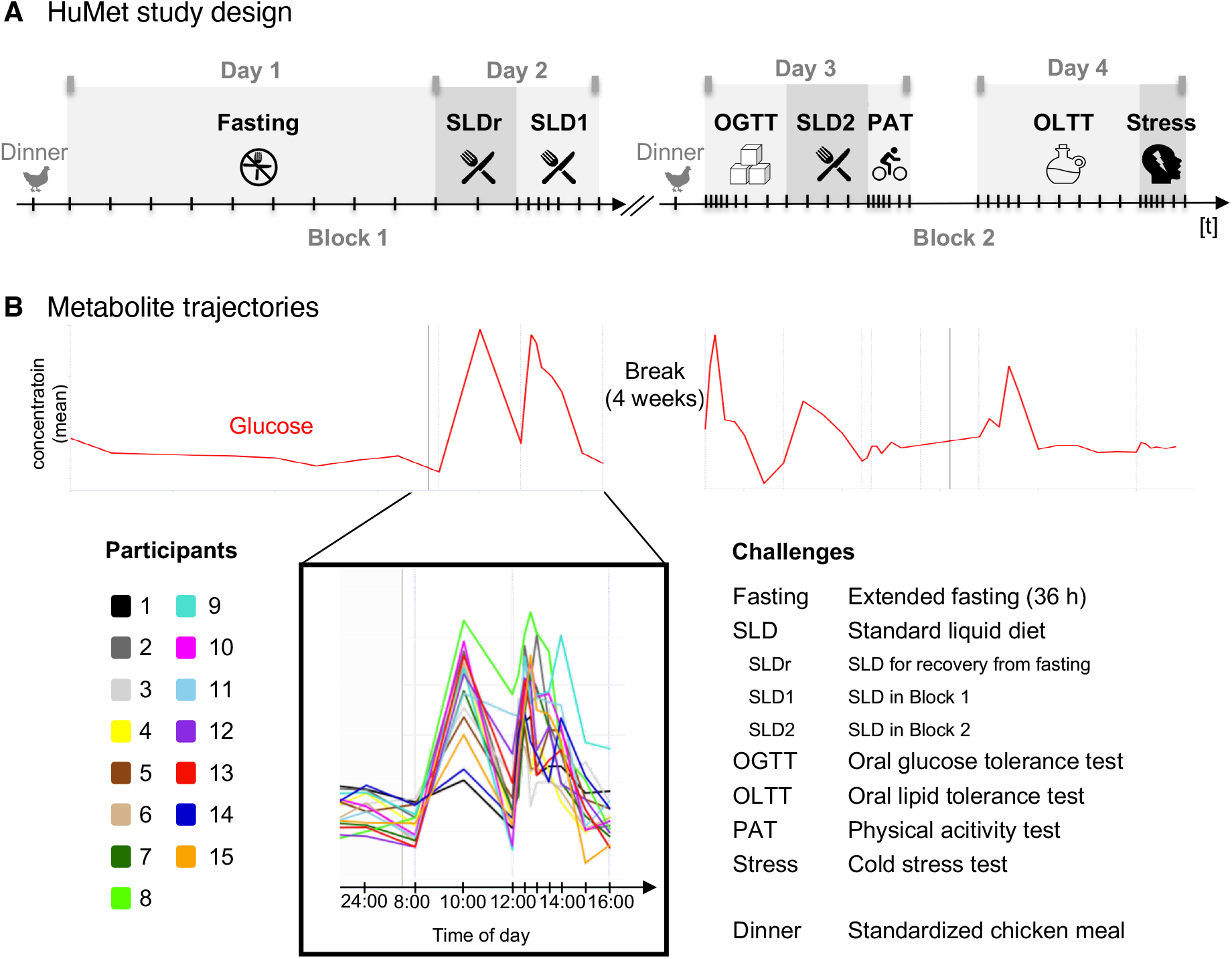
Metabolite profiles across six physiological challenges. (**A**) A sequence of metabolic challenges was applied over two study blocks, each covering a period of two days. All participants had the same chicken meal for dinner at 7 pm on the evening before each block. Five of the six challenge tests were applied once, while participants were exposed to the mixed meal challenge three times (SLDr, SLD1, SLD2). Challenge tests along with their abbreviations as used in the scheme are listed in B. Plasma, urine, and breath samples were collected at up to 56 time points in variable time intervals (15 min - 2 h) depending on the challenge. (**B**) In the repository, various representations of metabolite time courses can be visualized. Here, the time course of plasma glucose is shown as an example. The red line represents the mean levels over time, i.e., levels for the 15 study participants have been averaged at each of the 56 time points. The insert zooms into the 15 individual metabolite time courses for challenges SLDr and SLD1 (see legend), colored by study participants.

To identify metabolites whose abundances significantly changed in response to a challenge, we performed paired t-tests for each metabolite and time point during/after the challenge (within the time frames given in **Table 2**) compared to the challenge-specific baseline. After adjustment for multiple testing, this analysis yielded 620, 27, 117, 101, and 21 significant hits, comprising 220, 15, 66, 64, and 16 metabolites that changed at various time points during/after extended fasting, glucose/mixed/high-fat meals, and physical activity, respectively (**Supplementary Table 3**). Stress did not show any significant hit after correction for multiple testing (Bonferroni). For each of the challenges, **Table 2** lists those significantly altered metabolites that showed the lowest p-value and/or largest fold-change (decrease/increase) observed. As an example, the ketone body 3-hydroxybutyrate (BHBA) in urine showed the largest increase after 36 hours fasting when compared to overnight (12 h) fasting (log2 fold change (log2fc) 7.7). Significant increases of BHBA in plasma were observed before those in urine but only after prolonged fasting (log2fc 3.0 after 22 hours fasting), indicating the generation of ketone bodies for energy supply in this phase. Comparing the observed levels of plasma BHBA during extended fasting to those measured after the OLTT, we found similar levels of this ketone body 6 hours after ingestion of the lipid-rich challenge drink. All statistical results are provided in tabular form as well as in interactive volcano plots within the HuMet Repository (module *Statistics*).

**Table 2.**
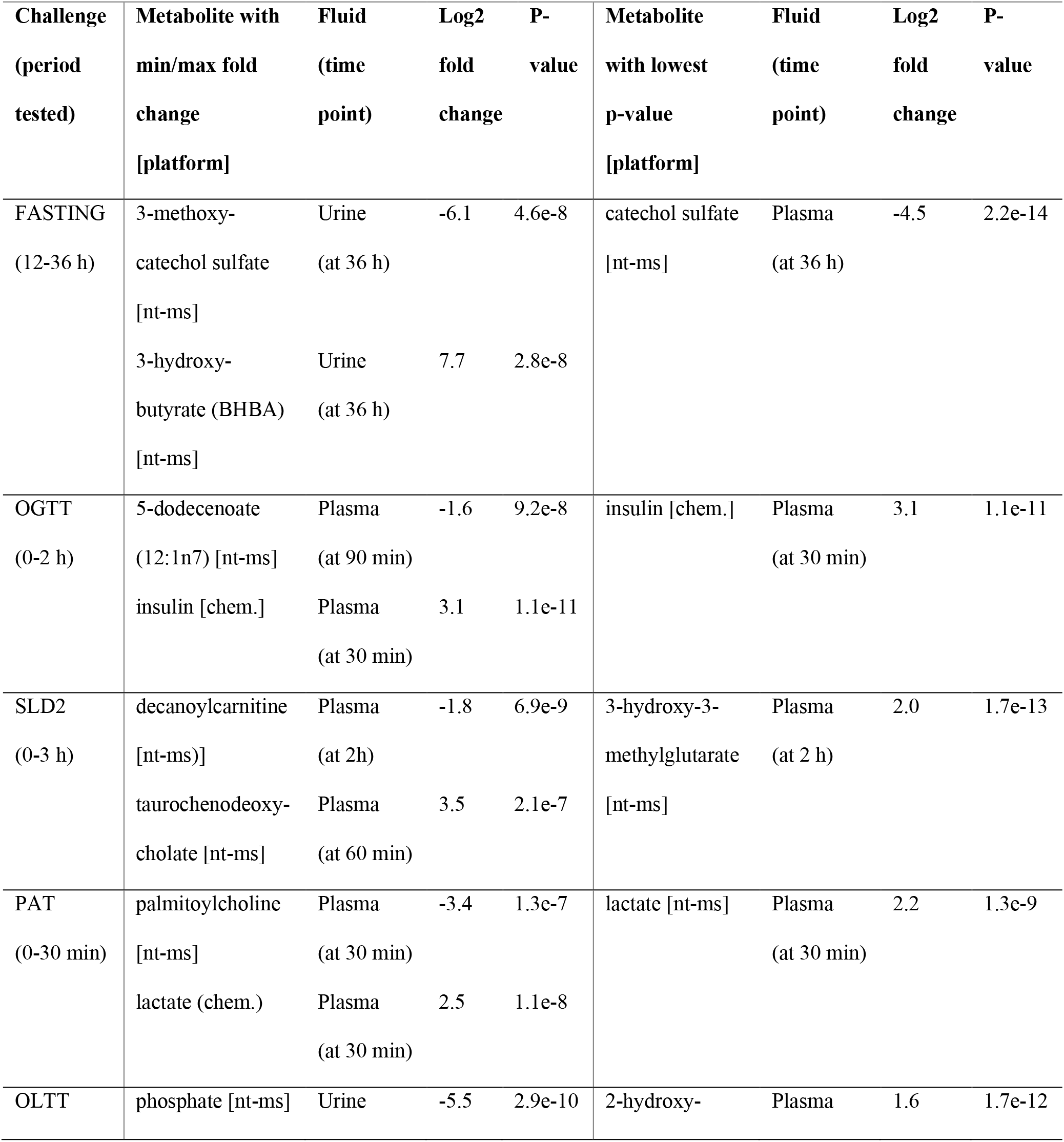
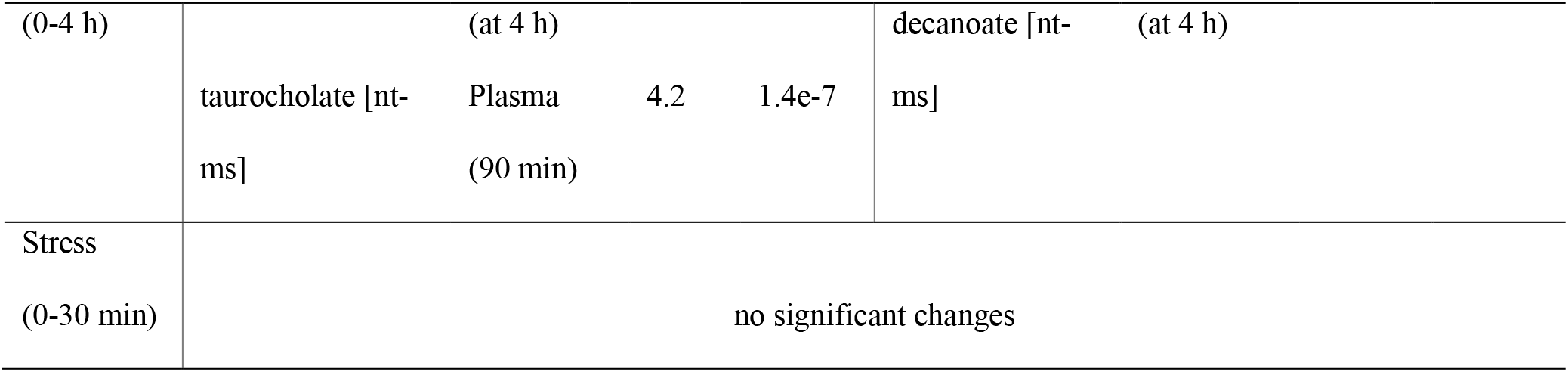
Metabolites with largest changes after each challenge.

### Data-derived metabolic networks provide molecular context for metabolite changes

To allow inspection of dynamic metabolic changes in the context of metabolic pathways and the overall metabolism, we generated different types of metabolic networks covering the metabolites of the targeted and non-targeted MS-based platforms in plasma and urine (Biocrates p150, Metabolon HD4), which were applied to samples from all participants and time points:

(i) *Knowledge-based* networks: These networks connect metabolites by their pathway membership based on existing pathway or metabolite class definitions (e.g., KEGG ^23^) as annotated by the providers of the metabolomics data; (ii) *Data-derived* networks: Here, we built on our previous findings that connecting metabolites based on their significant partial correlations in blood and urine reconstructs known metabolic networks from cross-sectional data, yielding so-called Gaussian graphical models (GGMs) ^24–26^. Applying a partial correlation method that takes the longitudinal design of data into account ^27^, we constructed GGMs based on the HuMet data from blood and urine for each fluid and platform separately (*single fluid networks*) (see **Methods**). For plasma, we additionally generated GGMs combining data from multiple platforms (Metabolon HD4, Biocrates p150, in-house biochemistry). For data from the Metabolon HD4 platform, we connected the plasma- and urine-specific GGMs into *multi-fluid networks* by linking the two nodes representing the same metabolite in each fluid by an additional edge. For example, the plasma network comprising metabolites from the Metabolon HD4 platform contains 339 edges connecting metabolites with partial correlations ≥ 0.12 (see **Methods** for details on cutoffs). Merging the plasma network with the network inferred from urine metabolites (Metabolon HD4 platform, partial correlations ≥ 0.09; see **Methods**), which consists of 227 edges, results in a multi-fluid network, where 333 edges connect the same metabolites measured in plasma and urine.

To test whether the GGMs inferred from longitudinal metabolomics data can also reconstruct metabolic networks, we assessed the known pathway distances of metabolites that were connected via an edge in the plasma GGM using the human pathway maps in KEGG ^23^. This analysis was possible for 74 out of the 339 edges, for which both metabolites were mappable to KEGG (n=129), and which were represented in a KEGG pathway of human metabolism. For 29 out of the 74 edges, the connected metabolites were also directly linked in KEGG (i.e., showed a pathway distance of 1); for 22 edges, we observed pathway distances of 2 or 3. A summary of results is provided in **Supplementary Table 4**. In a bootstrapping approach, in which we generated 1000 networks with the same topology but randomized node labelling, a maximum of 6 edges with pathway distance 1 was found in only one of the 1000 networks; we obtained similar results when performing the analogous analysis of the networks reconstructed from urine metabolites (**Supplementary Table 5**), confirming the applicability of GGMs for metabolic network reconstruction from the longitudinal HuMet data.

Using these data-derived metabolic networks, we mapped temporal changes in the abundances of metabolites by coloring nodes according to the metabolites’ log 2-fold changes during each challenge. This mapping allows for a holistic, metabolism-wide overview of time-resolved challenge responses (see also showcase *Systemic metabolic responses*).

### A web-based resource for data visualization and exploration

To facilitate access to data and results from the HuMet study, we set up the HuMet Repository as a web-based framework holding the complete HuMet data set and providing four modules (*Selection*, *Time course*, *Statistics*, and *Network*) for interactive data exploration (**Figure 1**).

The user can filter the dataset to form subsets for visualizations and analyses across these modules. Filtering includes restrictions on specific time points, challenges, subjects, metabolomics platforms, and sample types. The user can also choose between various data transformation options, including data scaling, imputation, and data representation as log2-fold changes. Plots generated as part of the different modules are interactive and can be downloaded, together with the data that have been used to generate the plots. Furthermore, the framework allows users to download the complete data as well as selected subsets and transformations thereof for further analysis.

The module *Selection* allows users to select from the complete set of 2,656 metabolites in the HuMet dataset. For selecting metabolites of interest, we provide interactive tables where metabolites can be sorted or filtered according to various assigned properties including metabolite classes, metabolomics platform, biofluid, and KEGG ^23^ or HMDB ^28^ identifiers. Moreover, metabolites can be selected by specifying pathways of interest using either pathways as defined by KEGG or by the metabolomics platform (‘annotated’). In addition, the user can search for metabolites that show a similar temporal profile as a specified reference metabolite (see also showcases *Prior Exposure* and *Platform Comparison*). Thereby, metabolites can be ranked by their similarity to the reference using different distance measures (Euclidean, Manhattan, Fréchet) or correlation (Pearson). To facilitate the comparison of different sets of metabolites, the user can assign metabolites to different groups (“bags”) and can toggle between them when using other modules.

In the module *Time course*, the user can visualize and compare temporal trajectories of selected metabolites over the four days of the HuMet study, comprising the six different physiological challenges. For each metabolite, the subject-specific temporal profiles are displayed in different colors, using the same color coding for participants consistently throughout the repository (**Figure 2B**). The visualization is adaptive to various data transformations such as z-scoring, imputation of missing values, and log2 fold changes related to challenge baseline. Mean time courses connecting the mean levels of participants at each time point can be displayed for multiple metabolites in one plot to facilitate visual exploration and comparison of temporal changes between metabolites.

For identifying metabolites whose abundances significantly changed in response to a challenge or between two user-defined time points, the HuMet Repository provides hypothesis testing within the *Statistics* module. Thereby, the user can choose between different approaches for multiple testing correction. For amenable exploration, all statistical results are provided in tabular form as well as in interactive volcano plots.

The module *Networks* allows users to inspect metabolites of interest in the context of metabolic pathways and reconstructed metabolic networks. To this end, we provide knowledge-based and data-derived networks for the metabolites from the two MS-based metabolomics platforms. For network generation based on the pre-calculated pairwise partial correlations between metabolites, the user can choose from different cutoffs, above which edges are drawn (using fixed partial correlation thresholds ^29^; at significance threshold corrected for multiple testing of edges (FDR, Bonferroni)), leading to various alternatives of single fluid networks for plasma and urine. To allow exploring the metabolic response to the different challenges within these networks, the user can map the log2 fold changes of metabolites (in relation to their challenge baselines) onto the metabolite nodes in the network using red (increase) and blue (decrease) color gradients. For the generation of the multi-fluid networks based on the new Metabolon HD4 plasma and urine datasets, we merged the corresponding single fluid networks by connecting the same metabolites measured in plasma and urine by an additional edge. Using this approach, changes in urine and plasma metabolites can be displayed in parallel.

### Showcase Prior Exposure: Identify metabolites with washout-like temporal profiles

In the first showcase, we seek to identify metabolites that originate from exposures such as foods or drugs, to which participants had no access during the 2 x 2 days of the study, using the HuMet Repository. Prior to each of the two blocks in the HuMet study (**Figure 2**), all participants ate the same “chicken with vegetables” meal (prepared from a packaged frozen instant meal), containing a complex mixture of dietary ingredients that were not included in any of the liquid meals provided during the 4-day study phase (e.g., meat or vegetables).

Methylhistidines have been suggested as biomarkers that reflect chicken meat intake ^30, 31^. In our participants, plasma levels of 3-methylhistidine exhibited a washout-like temporal profile with a steady decrease after chicken intake. The profiles showed minimal interference with stimuli during the study phase, which is a prerequisite for a true dietary biomarker (**Figure 3**). We therefore chose this metabolite as a starting point for the search of metabolites with similar kinetic characteristics, potentially indicating further prior exposures of the participants. We used the similarity search option in the module *Selection* to rank metabolites by the distances of their temporal profiles to the reference profile of 3-methylhistidine in plasma.

**Figure 3.**
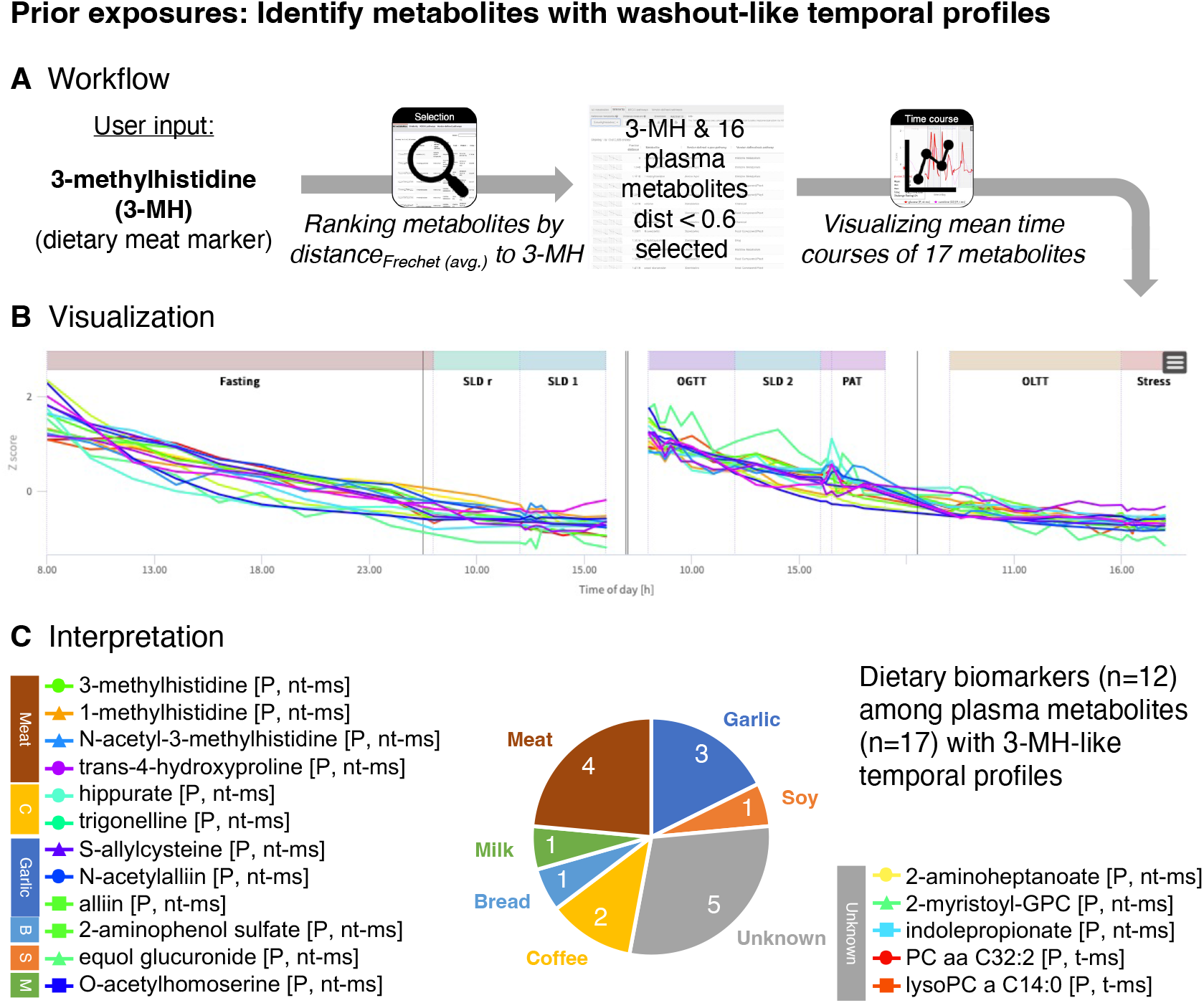
Exploration of metabolites from prior exposures. (**A**) Workflow to identify metabolites with similar trajectories as the reference plasma metabolite 3-methylhistidine (3-MH), a dietary marker for meat intake, using the similarity search implemented in the module *Selection*. (**B**) Time courses of 3-methylhistidine (3-MH) and the 16 plasma metabolites with most similar trajectories (Fréchet distance < 0.6) as visualized within the module *Time Course*. (**C**) Out of the 17 metabolites 12 are known biomarkers for various food items that have not been provided to participants during the study blocks, indicating exposures prior to the study.

We found S-allylcysteine in plasma to be the metabolite with the most similar temporal profile, showing a distance (Fréchet) of 0.2712 to plasma 3-methylhistidine; 34 additional metabolites had distances less than 0.6, with 16 being plasma metabolites. Out of these 16 plasma metabolites 12 are metabolites (or direct derivatives of metabolites) listed in FooDB ^32^ and/or are linked to food-related exposures in the Exposome Explorer ^33^. These 12 metabolites indicate putative exposures to meat, garlic, bread, coffee, milk, and soy (**Table 3; Supplementary Table 6)**. Most of these metabolites were detectable in almost all participants and at most time points. In contrast, equol glucuronide was only detected in two individuals (subjects 1 (6 time points of the first block) and 8 (all time points)), respectively. Equol is generated from daidzein, an isoflavone that is commonly found in legumes, particularly in soy. Only a fraction of the human population (∼50%) is able to convert daidzein into equol (which can then be further sulfated and glucuronated) ^34, 35^. The ability to produce equol presumably depends on the composition of a person’s microbiome and might be crucial for the health benefits that have been linked to soy isoflavones. At least two HuMet participants have this ability but only for one of them equol glucuronide was detected in both blocks of the study, suggesting that (out of the two) only this person was exposed to soy (or other daidzein-containing food) before each of the study blocks.

**Table 3.**
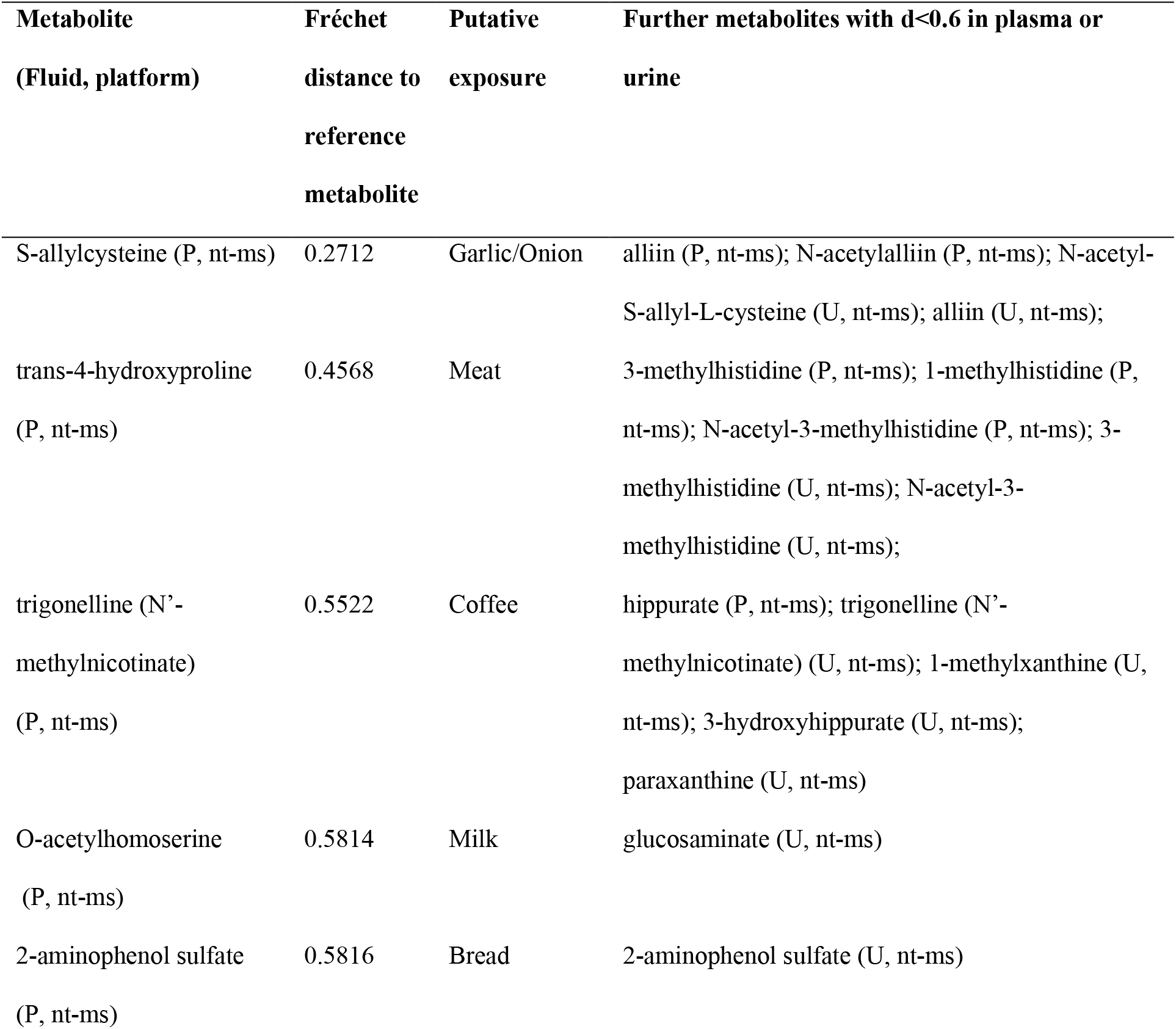
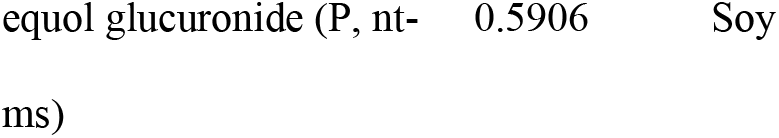
Food-related prior exposures linked to metabolites with similar washout-like temporal profiles as 3-methylhistidine. (dietary marker for (chicken) meat intake ^31^).

### Prior exposures: Identify metabolites with washout-like temporal profiles

Inspecting the list of metabolites with similar washout-like trajectories as 3-methylhistidine, we found five metabolites that have not been reported as dietary biomarkers. Interestingly, three out of these five metabolites are lipids that contain a C14 fatty acid residue (2-myristoyl-GPC (14:0), (dFréchet = 0.5415; PC aa C32:2 (mainly consisting of PC (14:0_18:2) ^22^), dFréchet = 0.5772; lysoPC a 14:0, dFréchet = 0.5909). The steady decline of these metabolites over the two blocks, when participants were only exposed to the highly standardized challenge drinks, suggests that dietary choices (but not acute fasting status or macronutrient composition of the challenge drinks) modulate the levels of these complex lipids.

Taken together, this use case demonstrates the value of the similarity search as implemented in the Module *Selection* to depict metabolites showing the same dynamic behavior.

### Showcase Platform comparison: Compare metabolites across metabolomics platforms

HuMet samples were profiled using a variety of metabolomics platforms, considering that no single approach can cover all parts of metabolism in sufficient quality ^36^. While their coverage of metabolites is mostly complementary, comparable measurements are available from the Metabolon HD4 platform and the Biocrates p150 kit for various amino acids, acylcarnitines, and glycerophospholipids. In this use case, we were interested to what extent measures for matching metabolites correlate between the platforms. This comparison is of particular interest for matching metabolite pairs where the platforms do not quantify the exact same analytes due to the different measurement techniques. As an example, the 43 matching metabolite pairs listed in Yet et al. ^37^ for Biocrates p150 (t-ms) and a prior version of Metabolon HD4 (nt-ms), include the pair H1 (Hexose) (t-ms)/glucose (nt-ms). While the non-targeted technique measures the (relative) abundances of glucose, the most abundant hexose in human blood ^38^, the targeted assay measures the concentrations of all hexoses as a sum.

Out of the 43 metabolite pairs, data on 38 pairs are available in the HuMet dataset. Overall, we observed a high correlation of measurements for the investigated metabolite pairs across the two platforms in HuMet, with a median correlation of 0.75 (**Figure 4**; **Supplementary Table 7**). Also, glucose (nt-ms) and H1 (hexose) (t-ms) measurements were highly correlated (r = 0.87), which is in line with measured plasma hexose consisting mostly of glucose in humans. Only four pairs showed comparably weak correlations (r < 0.5) (**Figure 4**). In particular, the acylcarnitine measures butyrylcarnitine (nt-ms)/C4 (butyrylcarnitine) (t-ms) (r = 0.18) and glutarylcarnitine (nt-ms)/C6-OH (C5-DC) (t-ms) (r = 0.18) showed differences between the platforms. In the first case, the reason for this difference could be that the Biocrates p150 measure labeled as C4 (butyrylcarnitine) includes the isobaric isobutyrylcarnitine, while these two metabolites are measured as two separate analytes on the Metabolon HD4 platform. Correlation analysis of metabolite isobutyrylcarnitine (nt-ms)/C4 (butyrylcarnitine) (t-ms) (r = 0.82) indicates that C4 (butyrylcarnitine) (t-ms) and/or its dynamic changes might indeed be dominated by isobutyrylcarnitine. This is of particular interest as butyrylcarnitine and isobutyrylcarnitine derive from two fundamentally different pathways linked to the degradation of fatty acids and to the degradation of branched-chain amino acid, respectively. A similar scenario can be assumed to underly the low correlations between glutarylcarnitine measured on Metabolon HD4 and the analyte labeled as C6-OH (C5-DC) measuring glutarylcarnitine and hydroxyhexanoylcarnitine together using the Biocrates p150 kit.

**Figure 4.**
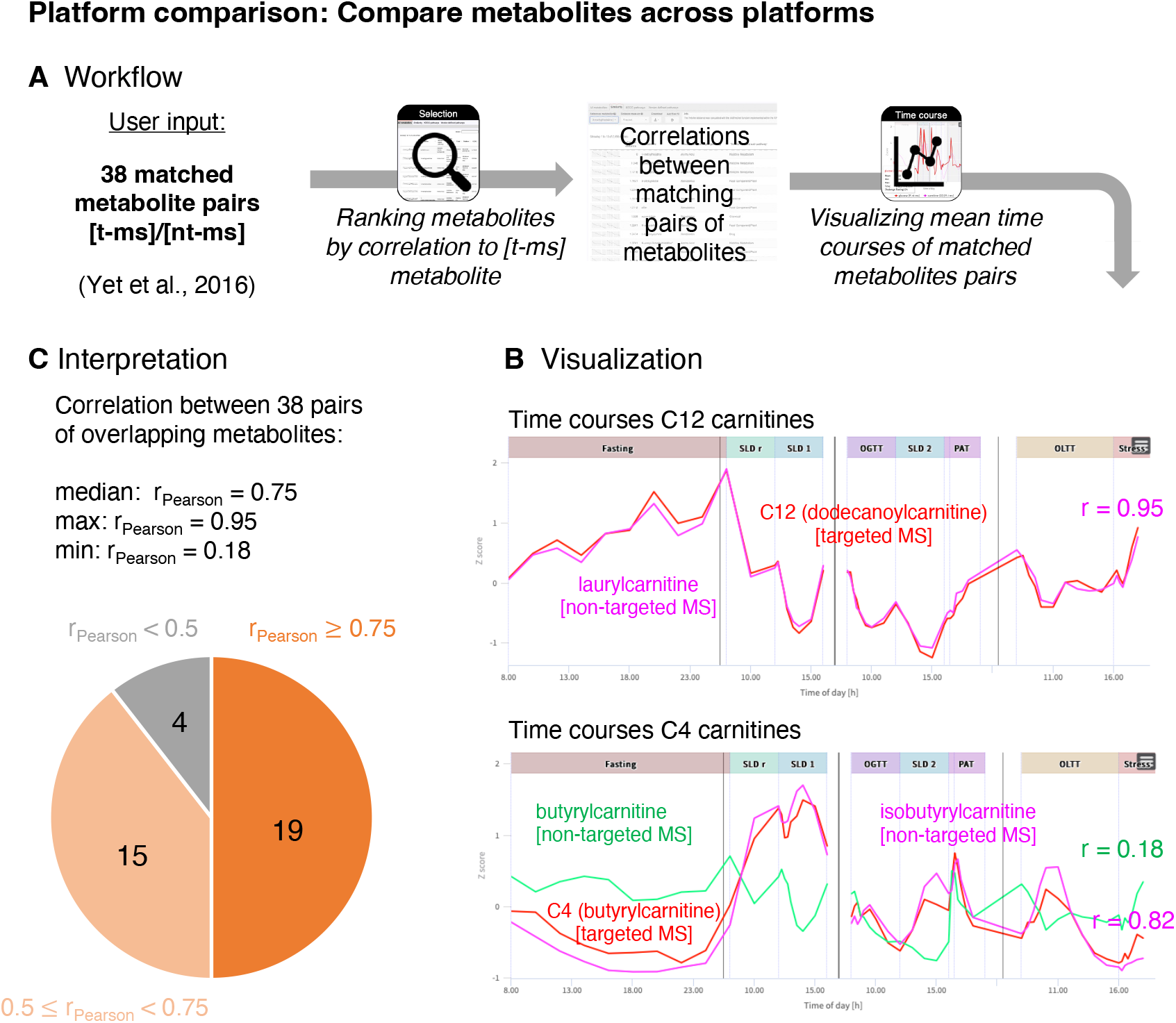
Comparison of measurements from different platforms. (**A**) Workflow to explore the concordance of measurements for 38 pairs of matching metabolites from the non-targeted (Metabolon HD4) and targeted (Biocrates p150) platform (pairs taken from Yet et al. ^37^). Pearson correlations of metabolites (across all time points and all subject’s individual metabolite curves) are provided through the similarity search implemented in the module Selection. (**B**) Trajectories of the pair with strongest correlation (laurylcarnitine (P, nt-ms)/C12 (dodecanoylcarnitine) (P, t-ms) with r = 0.95 and the pair with weakest correlation (butyrylcarnitine (P, nt-ms)/C4 (butyrylcarnitine) (P, t-ms)) with r = 0.18 are shown as displayed within the module *Time Course*. In case of the C4 carnitines, the measurements for the isobaric isobutyrylcarnitine (P, nt-ms), which is added to the time course plot, showed a much stronger correlation with the C4 measurement from the targeted platform, indicating that C4 (butyrylcarnitine) (P, t-ms) and/or its dynamic changes might be dominated by the isoform isobutyrylcarnitine. (**C**) Overall, the concordance of measurements from the two platforms is high with a median correlation of measurements of r = 0.75 and only four out of 38 pairs with correlations below 0.5.

Taken together, this use case demonstrates the value of the HuMet Repository for comparing measurements from different metabolomics platforms. Here, the availability of time-resolved metabolomics data from multiple platforms for the same participants combined with the exploration tools implemented in the repository (specifically the similarity search (*Selection* module) and aggregation of metabolite time courses plots (*Time Course* module)) facilitates the comparison and demonstrates how measurements from different platforms can inform each other.

### Showcase Systemic metabolic responses: Reveal and compare systemic responses to challenges

In this use case, we seek to answer the following questions: (i) *“Which areas of metabolism change after extended fasting compared to standardized overnight fasting in the reconstructed metabolic network?”* and (ii) *“How do metabolic responses in particular pathways compare between three different nutritional challenges?”*. Using the inferred metabolic networks within the repository’s *Networks* module, we can visualize and depict time-dependent responses to metabolic challenges in a metabolism-wide manner.

To get a global view of changes in metabolism after prolonged fasting, we chose the multi-fluid network (Metabolon HD4 plasma and urine) with default cutoffs for the underlying partial correlation (see **Methods**). On this backbone, we mapped statistical results comparing metabolite levels after extended fasting (36 h; time point (TP) 10) with levels after standardized overnight fasting (12 h; TP 1). In the resulting network, we saw widespread metabolic changes with prominent increases (indicated by red color with high saturation and large circle sizes of metabolite nodes) in various pathways, including a metabolite cluster containing ketone bodies (and their precursors from ketogenic amino acid degradation) and a cluster containing acylcarnitines (**Figure 5**). Also, increases in other clusters became apparent (indicated by circles in **Figure 5B**), for example clusters containing (i) sulfated bile acids (and steroids), in particular the monohydroxy bile acid derivative taurocholenate sulfate in blood and urine; (ii) nucleotides (xanthine, hypoxanthine) and metabolites of the citrate cycle (malate, fumarate), which also increased during exercise; and (iii) dicarboxylic fatty acids (mainly C10 – C18). Most decreases (blue color) were observed in clusters containing xenobiotic metabolites or metabolites that have been linked to the human gut microbiome (**Figure 5**).

**Figure 5.**
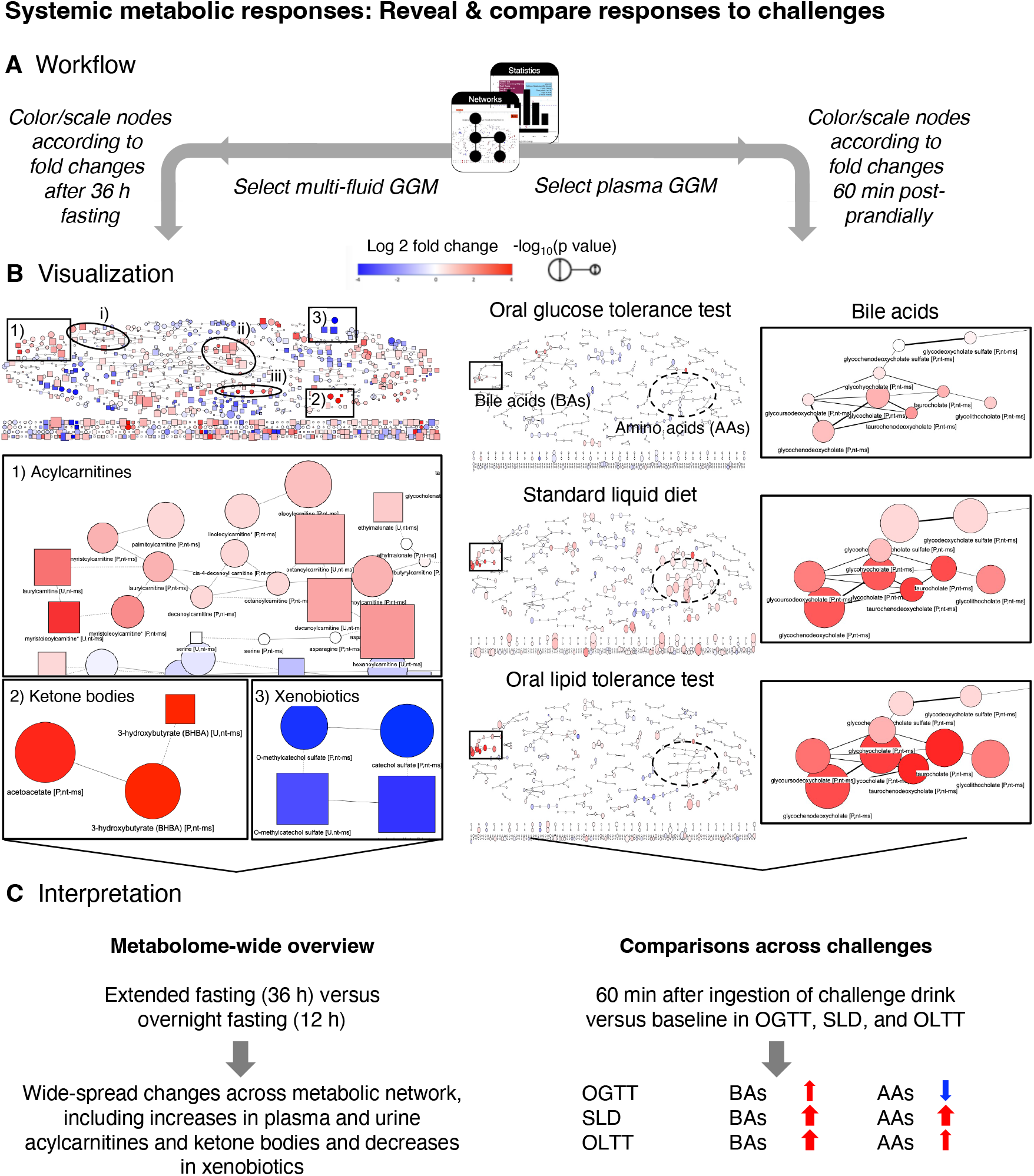
Contextualization of metabolic responses to challenges within reconstructed metabolic networks. (**A**) Workflow to explore metabolite responses to challenges from a holistic, metabolome- wide perspective. To get an overview of changes after extended fasting (36 h) compared to overnight fasting (12 h), we select the multi-fluid metabolic network derived from the non-targeted metabolomics data in plasma and urine provided in the module *Networks* (left). For comparison of metabolite changes within particular pathways (here: bile acids and amino acids) 60 min after ingestion of the challenge drink, we select the single fluid network generated based on plasma levels of the non-targeted (Metabolon HD4) and targeted (Biocrates p150) platform (right). To visualize responses, we map the log2 fold changes and p-values resulting from t-tests comparing metabolite levels after the respective challenge with the corresponding baseline levels (module *Statistics*). Color displays the log2 fold change between challenge baseline and chosen time point, with red indicating an increase in metabolite concentration and blue indicating a decrease in metabolite concentration. Node size depicts the -log10 p-value of metabolic changes between challenge baseline and chosen time point. Thereby, node size increases with lower p-value. (**B**) Coloring and scaling of metabolite nodes according to changes after extended fasting of 36 h (versus overnight fasting (12 h)) shows that various parts of human metabolism are affected by adaptations to this challenge (left). This includes pathways such as beta-oxidation of fatty acids (indicated by increasing acylcarnitine levels in blood and urine (see zoom-in for box 1) as well as the generation of ketone bodies (indicated by their increased levels in both fluids; see zoom-in for box 2). Moreover, increases in metabolite levels are observed in further clusters delineated through circles ((i) sulfated bile acids (and steroids); (ii) nucleotides (xanthine, hypoxanthine) and metabolites of the citrate cycle (malate, fumarate); (iii) dicarboxylic fatty acids (mainly C10 – C18); no zoom-ins provided). Decreases (blue color) are seen for various pathways of xenobiotic metabolites, including benzoate metabolites (see zoom-in for box 3). Coloring the network based on plasma metabolites by changes 60 min after ingestion of the challenge drinks (right) revealed a cluster of bile acids that similarly increased in SLD and OGTT and to a lesser extent also in response to OGTT. In contrast, a cluster containing most amino acids, showed decreases in OGTT and increases in SLD and OLTT, with less effect observed in the latter case (though 65% of the same protein mix was ingested with the OLTT drink as with the SLD drink). (**C**) Using the reconstructed single- and multi-fluid metabolic networks helps to get a metabolome-wide overview over dynamic metabolic changes in response to specific challenges and facilitates comparison of effects between different challenges.

For comparison of metabolic responses across challenges, we selected a single fluid GGM (Metabolon HD4 Plasma, Biocrates p150 plasma and in-house biochemistry, partial correlation (pcor) ≥ 0.12). Here, we mapped metabolic changes obtained for the OGTT (60 min vs. baseline), SLD (60 min vs. baseline) and OLTT (60 min vs. baseline) and focused on two modules that exhibited consistent and different changes across challenges, respectively (**Figure 5**): (i) a cluster containing different bile acids consistently increased after one hour in all three challenges and (ii) a cluster containing various amino acids that showed considerably different responses between challenges: the majority of metabolites within this cluster decreased 60 min after glucose ingestion in the OGTT, while they increased after ingestion of the SLD drink and less extensively also after ingestions of the lipid-rich OLTT challenge drink. More detailed inspection of the bile acid time courses using the *Statistics* module confirmed that, for five (glyco- and taurocholate, glyco- and taurochenodeoxycholate, and taurodeoxycholate) of the bile acids in the cluster, the observed increases were statistically significant not only one hour after ingestion of the lipid-containing SLD and OLTT challenge drinks but also one hour after ingestion of glucose in the OGTT with even higher fold changes observed after 15 and 30 min. Displaying the individual time courses of these bile acids across challenges in all participants using the *Time Course* module also showed that relative abundances and maximal fold changes of these bile acids strongly vary between individuals, challenge drinks, and even between different contexts: When the SLD challenge drink was ingested “for lunch” four hours after ingestion of the OGTT drink (SLD2), observed maximal log2 fold changes for the five bile acids were almost as high as in the OLTT (log2fc ∼ 3-4); the same maximal log2 fold changes were seen when the SLD drink was ingested in the morning at 8 am after the prolonged fasting (SLDr). In contrast, when the SLD drink was provided “for lunch” four hours after the first SLD drink in the morning on day two (SLD1), the maximal fold changes were smaller (log2fc ∼ 2; similar as the bile acid fold changes after OGTT), as the bile acid levels had not returned to the morning levels after these four hours.

Taken together, this use case demonstrates the usefulness of the HuMet Repository to explore and compare metabolic responses in the context of metabolic networks and across challenges and pathways.

## Discussion

The HuMet Repository described herein provides an easily accessible and explorable reference for typical metabolic responses to all-day challenges in healthy male individuals. The contained time-resolved metabolomics data set is unique regarding its metabolite coverage, with data from multiple different platforms, together capturing most areas of human metabolism in blood and urine. In particular, the non-targeted metabolomics data (plasma 595; urine 619), which we added to the data that already existed from the HuMet study, increased the breadth of metabolic pathways and enhanced the granularity of the metabolomic readout for exploring metabolism under challenge.

To bring the rich data set to the scientific community, we set up our repository in a way that goes beyond the idea of sharing data for re-analysis by experts according to the FAIR (Findable, Accessible, Interoperable and Reusable) principles: By offering interactive exploration and visualization tools, we enable users to query the data directly and flexibly while taking the burden of data handling for these complex, high dimensional data (15 subjects x 56 time/6 challenges x 2,656 metabolites/8 platforms x 4 biofluids) from them (e.g., by providing calculation of fold changes, imputation, biostatistics, mapping of metabolites onto metabolic networks). As a consequence, answering *ad hoc* questions like *“How robust is the metabolite that I identified as a disease biomarker in fasting versus non-fasting conditions?”, “How do dynamic changes in blood compare to those in urine for my metabolite of interest?”, “How individual is the metabolic response to physical activity among healthy subjects for my metabolite of interest?”, “Which metabolites exhibit the same longitudinal patterns over a challenge or the complete study duration as my metabolite of interest?”* is only a matter of clicks in the HuMet Repository, making the data thus more accessible for researchers with various backgrounds.

In addition to serving as a reference for questions regarding specific metabolites of interest, the HuMet Repository facilitates systematic explorations into metabolic responses across biochemical pathways, physiological challenges, and metabolomics platforms as we demonstrated in three showcases. Applying the built-in similarity search for extraction of metabolites with similar temporal profiles, we were able to identify metabolites with washout-like trajectories in our first showcase. In our second showcase, we demonstrated how assessing the similarity of metabolite trajectories can be used to compare readouts from different metabolomics platforms available for the same samples in the HuMet Repository. In contrast to previous work, where we used HuMet data for similar purposes ^22, 39^, we here performed data analyses solely using functionality of the HuMet Repository, without any additional effort for data processing, analysis, or visualization. In our third showcase, we highlighted the value of data-derived metabolic networks as provided within the repository to contextualize statistical results from metabolome-wide analyses and to compare them across six different challenges. In contrast to metabolic networks taken from knowledge-based maps such as KEGG, which typically omit many of the measured metabolites, the entire set of metabolites detected through the underlying metabolomics approaches is represented in data-derived networks. In the present study, we demonstrated that partial correlations calculated from the HuMet study, encompassing time-resolved data from only 15 individuals, yielded networks in which functionally related metabolites were grouped similarly as in networks derived from large cross-sectional data of more than 1000 individuals ^24–26^. The networks reconstructed based on HuMet data are, thus, adequate to provide metabolic context for the global exploration of challenge-induced temporal changes.

Besides demonstrating the general applicability of the HuMet Repository, our showcasing analyses also derived concrete new biological hypotheses: In our first case, we identified potential new markers of dietary intake by extracting metabolites that indicated exposures of our participants prior to the start of the two study blocks. Most of the metabolites that showed similar washout-like kinetic patterns as a known marker (3-methylhistidine) of a known prior exposure (meat) ^31^, were dietary biomarkers of further food items which were contained in the meal that was served to every participant at the evening prior to the study blocks or otherwise consumed before (e.g., metabolites of meat, garlic/onion, coffee, and soy). Interestingly, our analysis additionally revealed various phosphatidylcholines containing a C14:0 saturated fatty acid residue that showed the typical washout-like kinetic for most participants. These metabolites are not listed as dietary biomarkers in FoodDB ^32^ or Exposome Explorer ^33^ and are typically considered to be endogenous. Nonetheless, our results suggest that dietary choices strongly influence the blood levels of these metabolites. As bovine milk fat is rich in C14:0 ^40, 41^, the steady decrease of the C14:0 phosphatidylcholines within each of the two study blocks might reflect the “wash-out” of metabolites originating from cream, which was an ingredient of the served dinner. On the other hand, considering previous results from Altmaier et al., who reported an association of phosphatidylcholines with shorter fatty acid chains (< C20) and higher saturation with fiber intake ^42^, the observed effect of constantly decreasing levels of C14:0 phosphatidylcholines could also be related to the lack of fiber in the provided challenge drinks.

Results from our second showcase emphasize that the concordance of measurements from different metabolomics platforms for the “same” (matching) metabolites can vary depending on sampling time and conditions. On average, we saw high correlation of measurements across the investigated metabolites from the targeted Biocrates p150 and the non-targeted Metabolon platforms despite differences of what exactly is quantified for matching compounds between the two platforms by design (e.g., relative abundance of glucose (non-targeted) versus absolute concentration of all (isobaric) hexoses (targeted)). We thereby replicated results from previous cross-platform studies which indicated that these measures can be used largely interchangeably in cross-sectional data ^37, 43^. At a first glance, the measures labeled with ‘butyrylcarnitine’ were an exception: In the time-series data of the HuMet study, we only found weak correlation of the targeted ‘butyrylcarnitine’ analyte (representing the sum concentrations of isobaric C4 carnitines) with the non-targeted butyrylcarnitine measure. In contrast, under overnight fasting conditions in cross-sectional data of 1001 subjects, the correlation of the two measures was strong and the association of measured levels with a genetic variant in the ACADS locus, which encodes an enzyme converting butyrylcarnitine, was identified independent of the platform ^37^. However, in the time-series data of the HuMet study, we instead observed a strong correlation of the targeted C4 carnitine analyte (‘butyrylcarnitine’) with isobutyrylcarnitine measured as a separate analyte on the non-targeted Metabolon HD4 platform. Thus, our analysis revealed that, when dynamically monitoring targeted C4 carnitine concentrations over challenges, the concentrations resemble changes in isobutyrylcarnitine, which is linked to the degradation of branched-chain amino acids, rather than butyrylcarnitine, which is derived from the ß-oxidation of fatty acids. Hence, while the fasting C4 concentrations in the targeted analysis indeed reflected the inter-individual differences in butyrylcarnitine levels, this was not the case under non-fasting conditions.

When investigating the effects of prolonged fasting in our third showcase, where we used the repository’s network visualization and statistics functionality, we found that most measured metabolites in plasma and urine were affected by the fasting challenge to some extent. As expected from the lack of other energy sources, largest increases could be seen in metabolite clusters containing ketone bodies (and their precursors from degradation of ketogenic amino acids) and acylcarnitines, indicating burning of fat ^44, 45^. We also observed large increases in metabolite clusters containing dicarboxylic fatty acids (mainly C10 – C18), indicating fat degradation through peroxisomal fat oxidation. This process involves microsomal omega-oxidation and is known to occur during fasting ^46^, in particular when mitochondrial beta-oxidation is impaired as, for example, in specific rare monogenic diseases ^47^. The produced dicarboxylic acids have been suggested as regulators of beta-oxidation ^48^ with a potential role in hepatic lipid accumulation induced by fasting ^46^. Hepatic lipid accumulation and steatosis have not only been linked to starvation conditions but also to (chronic) excess of fatty acid influx into liver as in many cases of obesity and type 2 diabetes ^49^. While the HuMet participants were healthy and non-obese, we observed increases in these dicarboxylic acids also when there was an (acute) excess of fatty acids after ingestion of the high fat challenge drink for the oral lipid tolerance test (e.g., octadecanedioate log2fc: 1.45 (fasting), 1.02 (OLTT)), matching the hypothesis of dicarboxylic acids being mediators of lipid accumulation. Interestingly, in our study, we observed increases in dicarboxylic acid levels also after the physical activity test (e.g., octadecanedioate log2fc: 0.90 (PAT)) which might provide an explanation why prolonged physical exertion carries the risk of liver damage ^50^.

Another class of metabolites increasing during extended fasting comprised sulfated bile acid as well as sulfated steroid derivatives (e.g., taurocholenate sulfate, glycochenodeoxycholate sulfate, dehydroepiandrosterone sulfate). All these compounds are products of a sulfation reaction catalyzed by the enzyme sulfotransferase 2A1; common genetic variants in the encoding gene SULT2A1 have been reported to influence the blood levels of these compounds ^29, 51^. Despite various studies, the role of sulfotransferases in metabolic homeostasis is not fully understood yet and warrants further research ^52^.

While the showcases demonstrate the usefulness of the HuMet Repository, the data and our explorative approaches also have their limitations: First, the study was based on the idea to have a group of participants as similar as possible. This was realized with only 15 participants, all male, young, and normal weight with only minimal variation in BMI. This small sample size, however, clearly restricts the statistical power and leads to a lack of diversity in the study group, which, in turn, impedes the transferability of results to women or other age groups. On the other hand, the homogeneity of the group allows users to see inter-individual variation in metabolite levels and metabolic responses to the challenges in the absence of major sources that usually cause variation in metabolite levels, such as sex ^53, 54^, age ^55–57^, and BMI ^58^. Second, while the availability of data on six different challenges for the same participants facilitates comparisons across challenges, all participants were exposed to these challenges in the same preset order. Therefore, we cannot exclude carry-over effects between challenges. Nonetheless, the specific block-wise design of the HuMet study enabled for example the discovery of metabolites of prior exposure in the wash-out phase that are much more difficult or impossible to be found in studies that have only one time point or that performed only one challenge under standardized conditions. Third, the comprehensive coverage of metabolites mainly through non-targeted approaches comes with the downside that only relative abundances of metabolites are reported as opposed to absolute concentrations derived by the targeted methods. As a consequence, our resource cannot provide “normal concentration ranges” for the majority of measured metabolites, limiting the repository’s application as a quantitative reference. Also, as most data has been acquired from commercially available metabolomics platforms, we do not have access to the raw spectra for each measurement to share them publicly. However, for various large epidemiological cohorts, cross-sectional datasets are available from the same metabolomics platforms, which enables direct cross-links of HuMet results to results from metabolome- and genome-wide association studies ^29, 59–62^. Fourth, as we tailored the application to the specific HuMet study data, our repository is a standalone resource and not integrated into a larger metabolomics repository such as MetaboLights. However, the underlying non-targeted data is available via MetaboLights (MTBLS89). Finally, explorative data analysis as supported by our repository can only be used for generation of hypothesis, which need to be followed up by more specific and sophisticated data analyses and subsequent experiments.

## Conclusion

In conclusion, the HuMet Repository opens avenues for researchers with different backgrounds to explore human metabolism under challenge conditions. With its comprehensive coverage of the human metabolome in plasma and urine, its time-resolved metabolite profiles after six different metabolic challenges, and its interactive analysis and visualization tools, this repository allows for addressing numerous questions related to the dynamic changes in metabolism. Our showcases exemplify how the project-tailored webtool facilitates explorative as well as systematic data analysis e.g., to identify dynamic metabolic responses in a metabolome- and “challenge-wide” fashion, or to identify metabolites that exhibit synchronous trajectories.

While we here highlighted results from specific analyses and focused on the generated hypotheses, leveraging the HuMet Repository, users can tackle further question without the need of processing the data by themselves, e.g., *How much does the coupling of kinetic behavior across metabolites differ when comparing responses between different challenges (e.g., extended fasting and exercise)?*, and many more options e.g., for cross-linking metabolite measures from different analytical platforms or estimating the stability of metabolite levels within individuals. Moreover, as we used highly standardized challenges for testing lipid, fasting, or exercise tolerance, they can be repeated in future studies, for instance, in studies involving patient cohorts. Derived patient data could be directly compared to data in our repository for identifying deviations from the ‘normal’, healthy response. Therefore, the HuMet Repository could help unlock the full potential of standardized challenge tests and their metabolic readouts to identify metabolic aberrations, when they are not yet visible in the rested, unperturbed state, thereby enabling new concepts for disease prevention or early diagnosis.

## Methods

### HuMet study population and design

The present work is based on samples of the Human Metabolome (HuMet) study conducted at the Human Study Center of the Else-Kröner-Fresenius Center of Nutritional Medicine at the Technical University Munich. All details on study design, population, and existing data have been described previously ^11^. Briefly, 15 healthy male participants were recruited for the study. Participants were young (mean age of 27.8 years ± 2.9), had normal weight (mean body mass index (BMI) of 23.1 kg/m^2^ ± 1.8), did not take any medication, and did not show any metabolic abnormalities (**Supplementary Table 1**).

All participants underwent a series of six unique metabolic challenges within two 2-day test blocks (**Figure 1**). Twenty-four hours prior to each test block, participants were asked not to consume alcohol or engage in strenuous physical exercise. Participants were provided with the same meal (standard size, chicken-based with vegetables (FRoSTA Tiefkühlkost GmbH, Hamburg, Germany)) at 7 pm one day prior to each test block. During each study block, participants stayed within the study unit to reduce perturbation by environmental influences. Samples were collected at up to 56 time points in different intervals (every 15-240 min) over the study days depending on the collected biofluid (plasma, spot urine, exhaled breath condensate samples, breath air) and the particular challenge (**Supplementary Table 8**).

Challenges covered extended fasting, ingestion of three different drinks with unique macronutrient compositions, a physical activity, and a stress test: (i) The fasting challenge consisted of a 36-hour fasting period (from the dinner before block one until 8 am on day two in the first block). During the challenge, participants drank 2.7 liters of mineral water based on a defined drinking schedule. (ii) A standard liquid diet (SLD) drink was ingested at three occasions: SLDr – for “breakfast” on day two to recover from extended fasting, SLD1 – for “lunch” on day two, and SLD2 – for “lunch” on day three in the second block. The SLD drink consisted of a defined fiber-free formula drink (Fresubin® Energy Drink Chocolate, Fresenius Kabi, Bad Homburg, Germany), providing one-third of the daily energy requirement of each participant. (iii) The oral glucose tolerance test (OGTT) on day three (block two) consisted of a 300 ml solution with mono- and oligosaccharides, equivalent to 75 g glucose after enzymatic cleavage (Dextro O.G.T., Roche Diagnostics, Mannheim, Germany). (iv) The oral lipid tolerance test (OLTT) on day four combined two parts of the SLD and one part of a fat emulsion containing predefined long-chain triglycerides (Calogen®, Nutricia, Zoetemeer, Netherlands), while adjusting volumes per participant to provide 35 g fat/m^2^ body surface area. All challenge drinks were served at room temperature for ingestion within 5 minutes. (v) For the physical activity test (PAT) participants performed a 30 min bicycle ergometer training at a power level corresponding to their individual anaerobic threshold. (vi) In the cold stress test, participants were triggered by immersing one hand, up to wrist level for a maximum of 3 min in ice water. For a complete protocol of the challenge procedure and the collection of samples, see Krug et al. ^11^.

The ethical committee of the Technische Universität München approved the HuMet study protocol (#2087/08), which is in correspondence with the Declaration of Helsinki.

### Existing HuMet metabolomics data

The HuMet study samples were previously profiled on three different “in-house” and three different vendor-based platforms. Resulting metabolomics data were used as published and provided in Krug et al. ^11^ for integration into our HuMet Repository.

Measured metabolites from these six platforms and the analytical methods used were described in detail in Krug et al. ^11^ and are only briefly summarized here: (i) “*In-house biochemistry*”: Standard biochemistry assays were used to assess blood levels of glucose, lactate, insulin, and non-esterified fatty acids (NEFA) in 840 plasma samples (15 subjects x 56 time points). Venous plasma glucose and lactate concentrations were profiled using an enzymatic amperometric technique, insulin was measured by ELISA, NEFA were quantified in plasma by an enzymatic colorimetric method. All assays were performed at the Technische Universität München. (ii) “*In-house FTICR-MS”*: Flow injection electrospray ionization ion cyclotron resonance Fourier transform mass spectrometry (FTICR-MS) measurements were performed at Helmholtz Zentrum München. A total of 201 mass spectral features from volatile compounds were reported in 55 breath condensates samples (5 subjects x 11 time points of the first block). (iii) *“In-house PTR-MS”*: Proton transfer reaction mass spectrometry (PTR-MS) was used to profile 341 breath air samples (11 subjects x 31 time points). Analyses were performed by researchers from Helmholtz Zentrum München and yielded 106 mass spectral features of volatile compounds. (iv) *“Biocrates p150”*: Absolute*IDQ* p150 kits from Biocrates Life sciences AG, Innsbruck, were used to perform flow injection analysis mass spectrometry (FIA-MS) of 840 plasma samples (15 subjects x 56 time points), yielding quantities for 132 blood metabolites after quality control. (v) *“numares”/* “*Chenomx*”: NMR spectra of 810 plasma samples (15 subjects x 54 time points) and 195 urine samples (15 subjects x 13 time points) were determined by numares (formerly LipoFit Analytic GmbH, Regensburg, Germany). For plasma samples, a total of 28 metabolites were identified by the company based on these spectra. For urine samples, only the levels of six metabolites were extracted from the spectra (at Helmholtz Zentrum München) using the software Chenomx NMR suite 7.0.

All data were used as preprocessed and provided in Krug et al. ^11^, unless stated otherwise in the following.

### Non-targeted metabolomic profiling (Metabolon HD4)

In addition to the previously published metabolomics data, we acquired new data by profiling plasma and urine samples of the HuMet study on the non-targeted platform Metabolon HD4 using liquid chromatography coupled to mass spectrometry (LC-MS) at Metabolon, Inc. (Durham, NC, USA). This platform applies four different analytical methods optimized for measuring metabolites with different physicochemical properties: (i) a reverse phase (RP)/ultra-high-performance liquid chromatography (UPLC)-MS/MS method with electrospray ionization (ESI) in positive mode optimized for hydrophilic compounds, (ii) a RP/UPLC-MS/MS with ESI in positive mode optimized for more hydrophobic compounds (iii) a RP/UPLC-MS/MS with ESI in negative mode, and (iv) a hydrophilic interaction liquid chromatography (HILIC)/UPLC-MS/MS with ESI in negative mode. All methods utilized a Waters ACQUITY UPLC and a Thermo Scientific Q-Exactive high resolution/accurate mass spectrometer (operating at a mass resolution (m/Δm) 35,000) interfaced with a heated electrospray ionization (HESI-II) source. For methods i – iii, a C18 column from Waters (UPLC BEH C18-2.1x100 mm, 1.7 µm) was used, method iv utilized a HILIC column (Waters UPLC BEH Amide 2.1x150 mm, 1.7 µm).

Sample processing for and analytical procedures of the Metabolon HD4 platform have been described in detail previously ^20^. Briefly, EDTA-plasma and spot urine samples, which were kept at -80°C until analysis, were first thawed. Then, several recovery standards, which were carefully chosen not to interfere with the measurement of endogenous compounds, were spiked into 100 μl of every sample to allow chromatographic alignment and to monitor instrument performance. For protein precipitation and metabolite extraction, samples were mixed with methanol under vigorous shaking for 2 min (Glen Mills GenoGrinder 2000). After centrifugation, the resulting extracts were split into five portions for each sample: four aliquots for analysis by the different LC-MS methods and one aliquot for backup. The extracts were placed briefly on a TurboVap® (Zymark) to remove the organic solvent and then stored overnight under nitrogen. Before LC-MS analysis, the extracts were reconstituted in solvents compatible for the MS methods (with each reconstitution solvent containing a series of standards at fixed concentrations to ensure injection and chromatographic consistency). All described sample processing steps were automated using a MicroLab STAR® system from Hamilton Company (Reno, NV, USA).

For LC-MS analysis by method i (acidic positive ion conditions), the extracts were gradient eluted from a C18 column (see above) using water and methanol, containing 0.05% perfluoropentanoic acid (PFPA) and 0.1% formic acid (FA). For analysis by method ii (acidic positive ion conditions), the extracts were gradient eluted from the same C18 column using methanol, acetonitrile, and water, containing 0.05% PFPA and 0.01% FA. For analysis by method iii (basic negative ion conditions), the extracts were gradient eluted from a separate C18 column using methanol and water with 6.5 mM ammonium bicarbonate at pH 8. For analysis by method iv (basic negative ion conditions), the extracts were gradient eluted from a HILIC column using a gradient consisting of water and acetonitrile with 10 mM ammonium formate at pH 10.8. The MS analysis alternated between full scans (covering 70-1000 m/z) and data- dependent MS^n^ scans using dynamic exclusion.

Peak identification and alignment from the recorded spectra, were performed using Metabolon’s in-house hardware and software. Metabolites were identified by comparison of the experimental spectra to entries in Metabolon’s in-house library, which was collected from the measurement of commercially available purified standards (∼3,300 at time of analysis) or recurrent spectra from either named compounds (or classes), for which no authenticated standard was available (marked by a tag next to the metabolite name in **Supplementary Table 2**), or from structurally unnamed biochemicals. Note that, for the present study, only the results from measuring named metabolites were purchased from Metabolon. The area-under-the-curve (AUC) of the peaks indicated as the *quantification ions* in the library entries were used to quantify metabolites. To account for differences in solute concentrations, raw peak AUC values of metabolite in urine were normalized by osmolality. Raw peak AUC values (plasma) and osmolality-normalized peak AUC values (urine) of each metabolite were additionally normalized to account for instrument inter-day tuning differences by dividing the values of each metabolite at each run day by the median of values for the metabolite on this day (i.e., setting the run day medians to one). Before data release, a series of manual curation procedures were carried out at Metabolon to remove metabolite signals representing system artifacts, mis- assignments, and background noise and to confirm the consistency of peak identification and quantification among the various samples. This work was based on proprietary visualization and interpretation software.

As the focus of the HuMet study was on the dynamic changes of metabolite levels within individuals, samples from the same individual were measured on the same run day (plate) to the extent possible, leading to a run day design where the plasma samples of two participants were analyzed on three different run days while assigning samples of block 1 (days 1 and 2), samples of day 3 (block 2), and samples of day 4 (block 2) to the same run day, respectively. Within run days the order of samples was randomized. Due to the lower number of urine samples, the samples from all time points of two participants were measured on the same run day. Plasma and urine samples of subject 4 were measured in duplicates; for this subject, we used the mean of both measurements. Several quality control (QC) samples, which underwent the same sample processing as the HuMet samples, were measured spaced evenly among the experimental samples: Ultra-pure water samples served as process blanks; pooled matrix samples (CMTRX) generated from all HuMet samples (only for urine samples) and aliquots of a pool of well-characterized human plasma (MTRX4) (both for plasma and urine runs) served as technical replicates to assess process variability across run days of the analysis. Relative standard deviation of CMTRX (urine) and MTRX4 (plasma) measurements are provided in **Supplementary Table 2**.

As a result of the analyses of 833 plasma and 240 urine samples on the Metabolon HD4 platform, relative abundances (normalized peak AUCs) are available for in total of 595 plasma and 619 urine metabolites. These metabolites were assigned to eight chemical classes termed super-pathways (amino acids, carbohydrates, cofactors and vitamins, energy, lipids, nucleotides, peptides, xenobiotics), each being divided into two or more sub-pathways, resulting in a total of 78 and 68 sub-pathways for the plasma and urine metabolites, respectively (**Supplementary Table 2)**.

### Lipidyzer

Lipid concentrations in HuMet plasma samples of four participants were analyzed on the Lipidyzer^TM^ platform of AB Sciex Pte. Ltd. (Framingham, MA, USA) by Metabolon Inc., Durham, NC, USA. Samples were kept at -80°C until analysis. The protocol of lipid quantification using this platform has been described in detail elsewhere ^22^. In brief, after thawing, lipids were extracted from the plasma samples with dichloromethane and methanol following a modified Bligh-Dyer extraction. For analysis, the lower, organic phase, which included internal standards, was used and concentrated under nitrogen. Extracts were reconstituted with 0.25 ml of dichloromethane:methanol (50:50) containing 10 mM ammonium acetate and placed in vials for infusion-MS analysis on a Sciex 5500 QTRAP equipped with a SelexION^TM^ differential ion mobility spectrometry (DMS) cell, which allows separation of different (lyso)phospholipids [(lyso)phosphatidylcholines ((L)PCs), -ethanolamines ((L)PEs), -inositols (PIs)] and sphingomyelins (SMs). Extracts were analyzed using multiple reaction monitoring (MRM) in two sequential flow injection analysis (FIA) runs, alternating between positive and negative polarity. Free fatty acids (FFAs), tri- and diacyglycerols (TAGs, DAGs), ceramides (CERs), lactosyl-, hexosyl-, and dihydroceramides (LCERs, HCERs, DCERs), and cholesterylesters (CEs) were measured using separation through the DMS cell. Lipids were quantified relative to appropriate stable isotope labeled internal standards. Concentrations are provided in μmol/l. The Lipidyzer platform allowed for absolute quantification of 965 lipids distributed over 14 lipid classes: (CE, TAG, DAG, FFA, PC, PE, PI, LPC, LPE, SM, CER, HCER, LCER, DCER).

### Data preprocessing and transformations

Quality controlled and normalized existing and new metabolomics data were forwarded to integration into the HuMet Repository. Thereby, metabolite names and abbreviations were kept as provided by the specific platforms. Metabolite identifiers within the repository contain the platform specific name, information on the fluid, in which they were measured (P: plasma; U: urine; BA: breath air; BC: breath condensate), and information on the platform (nt-ms: Metabolon HD4; t-ms: Biocrates p150; Lipidyzer: Lipidyzer^TM^; NMR: numares/Chenomx; PTRMS: In-house PTR-MS; ICR: In-house FTICR-MS; chem.: In-house biochemistry). Named metabolites were assigned to the eight different metabolite classes (“super-pathways”) as used for the Metabolon HD4 platform and to “sub-pathways” according to the categories given by the platforms. We manually annotated metabolites with links to compounds in knowledge-based platforms, including KEGG, PubChem, and HMDB.

For samples, information on the fluid, the subject (1-15), and the time point (1-56) (**Supplementary Table 8**) are used for identification (some breath air measurement were between two time points as defined for plasma/urine; they are denoted by 10.5, 11.5, 27.5, 39.5; for the six NMR urine metabolites (ChenomX), a sample from an additional time point (57; day 4: 7 pm) was measured).

The following preprocessing steps and transformations were applied to all metabolomics data:

*Manual curation.* To identify and remove outliers/implausible values, we systematically filtered single data points whose log2-transformed values were outside the mean ± 4 times the standard deviation window for the particular metabolite and time point, while omitting data points from measurements within the first 30 minutes of a study challenge (to avoid deletion of biologically meaningful challenged-induced concentration peaks of subjects). As a result, we identified 163 outlier data points, of which 92 data points were excluded after manual inspection. This cleaned data set is integrated into our repository and can be downloaded from the website.

*Data transformations.* In addition to the original concentration or relative abundance values, we provide the data after further transformations for display in the *Time Course* module: (i) *z-scores* based on the log2-transformed concentrations/relative abundances to facilitate comparisons across metabolites and platforms, (ii) *log2 fold changes (block)* calculated between the time points within each block relative to the first time point of the respective block, and (iii) *log2 fold changes (challenge)* calculated between the time points in a specific challenge and the challenge baseline (see **Supplementary Table 8**).

*Imputation.* Some of the downstream statistical analyses used in the HuMet Repository, such as network inference with GGM, require a full dataset without missing values. Before imputation, the manually curated dataset was filtered for metabolites with less than 30% missingness (n=493) across all samples measured on the particular platform (**Supplementary Table 2**). Based on the filtered data set, we imputed missing values using the machine learning algorithm *missForest* (ntree=1500, mtry=22), which is implemented in the R package *missForest* (version 1.4). The algorithm is based on a random forest approach and imputes missing values by iteratively (maximum iterations = 10) predicting missing values using the available data ^63^. This allows for accurate imputation of missing data while preserving the underlying non-linear data structure ^64^.

### Statistical analysis/functionality

*Metabolite time course similarity*. We provide several distance measures (Fréchet, Euclidean, Manhattan) and Pearson correlation to rank metabolites according to their similarity in temporal profiles. All measures are calculated based on z-scored data and depend on user-selected settings such as the choices of subjects and time-range. The distance/correlation between the temporal curves of two metabolites is calculated within each subject first; subsequently, we calculate the average distance/correlation across all chosen subjects. We additionally provide Fréchet distance and Pearson correlation calculated based on the mean metabolite trajectories (mean z-score over all participants at each time point). The Fréchet distance (on average trajectories) is set to default within the similarity tool. It uses a window approach to search for the smallest distance between curves in a defined timeframe. This time frame is defined as follows: Maximum of +/- 30 minutes within all challenges except the extended fasting. Within the extended fasting challenge, we allow for comparison of time-points within a range of +/- 120 minutes.

We used the *dist* function implemented within the R package *proxy* (version 0.4-23) to calculate the Euclidean and Manhattan distances. The R package *stats* (version 3.6.2.) was used to calculate the Pearson correlation. To calculate the Fréchet distance we used the *distFrechet* function implemented within the R package *longitudinalData* (version 2.4.1.).

*Paired t-tests.* We use paired *t*-tests to test for significant changes in metabolite levels between two time points based on the log2-transformed imputed or non-imputed (selectable by the user) concentrations/relative abundances, using the function *t.test* implemented in the R package *stats* (version 4.2.3). To adjust for multiple testing, we offer corrections based on the false discovery rate (FDR) (q < 0.05) or Bonferroni (p < 0.05/(n metabolites * n time points)). The levels of adjustment are reactive to the number of metabolites and time points submitted to statistical analysis. The user can select the time range and the option whether only the last time point or all time points within the range are compared to the first time point. Results are visualized within a volcano plot by using the function *plot_ly* of the R package *plotly* (version 4.9.1). Each data point within the volcano plot can be colored by super-pathway or metabolomics platforms.

### Network generation

Knowledge-based networks were constructed based on the annotated super- and sub-pathway structure of metabolites. This structure provides a quick overview of available metabolites from different platforms.

Network inference of Gaussian Graphical Models (GGMs) is based on partial correlations of metabolite concentrations/abundances (single fluid, imputed and log2-transformed data). These models have previously demonstrated to reconstruct biological pathways from cross-sectional metabolomics data derived from Biocrates and Metabolon platforms ^24^. To calculate partial correlations for the HuMet data sets we used the shrinkage estimator approach “GeneNet”, which is available within the R package *GeneNet* (version 1.2.14), choosing the “dynamic” method for estimation. This method relies on the function *dyn.pcor* implemented within the R package *longitudinal* (version 1.1.12), which takes the longitudinal data structure with repeated measurements from the same participant into account ^27^. If both dynamic partial correlation and Pearson correlation between two metabolites were statistically significant at a 5% significance threshold, pairwise metabolite connections were integrated into the network. Thereby, the user can choose between Bonferroni or FDR correction for multiple testing or restrict edges in the displayed network to those greater than several pre-defined dynamic partial correlation values.

Using this approach, we inferred and provide multiple single fluid networks based on one or more plasma or urine datasets from different platforms. For the generation of the multi-fluid network based on the plasma and urine datasets from the Metabolon HD4 platform, we merged the corresponding single fluid networks by connecting the same metabolites measured in plasma and urine by an additional edge, closely following the procedures reported in Do et al. for creating an overlaid network ^25^.

### Implementation of the web-based repository

The HuMet Repository is written in R ^65^ using shiny, an R package that enables setting up web-based graphical user interfaces (GUIs) while allowing to execute R code on the backend. All R Packages used for building the interactive HuMet Repository are listed in **Table 4** and are categorized into general packages for GUI implementation, statistical analysis, data visualization and performance.

**Table 4.** R Packages used within the HuMet Repository.

| Functionality | Description | Package name (version) |
| --- | --- | --- |
| Basic shiny | Shiny repository app structure | Shiny (1.7.4) |
| Graphical user interface extension packages | General shiny extensions | shinydashboard (0.7.2)<br>shinydashboardPlus (2.0.3)<br>shinyWidgets (0.5.0) |
|  | CSS extensions | shinyBS (0.61)<br>shinycssloaders (0.3) |
|  | Shiny JS extensions | shinyjs (1.1) |
| Statistical analysis | Metabolite rankings | Stats (3.6.2)<br>longitudinalData (2.4.1.) |
| Data visualization | Table visualization | DT (0.27)<br>tableHTML (2.0.0) |
|  | Temporal plots visualization | Highcharter (0.7.0) |
|  | Network visualization | visNetwork (2.0.9)<br>igraph (1.4.1)<br>colorRamps (2.3) |
|  | Statistical plots | plotly (4.10.1) |
| Data processing |  | doParallel (1.9.15)<br>readxl (1.4.2)<br>writexl (1.4.2) |
| Individualization | Individual scripts | Collected with tag: rep_ ... |

The repository loads the preprocessed data, metabolite information, sample information upon session start. Thereafter, the repository is reactive to the user’s choices of options. These include exclusion of data points due to selected time points, subjects, and platforms. The repository visualizes the chosen data in interactive plots that, e.g., provide additional information via hover-over functionality, allow for zooming, and data-dependent coloring of data points.

## Acknowledgement

We thank all people involved in the planning and conduction of the HuMet study, in particular Susanne Krug, and all 15 volunteers who agreed to go through the four days of strenuous six challenges. We also thank Maria Littmann, who contributed to the code of an early prototype of the web application during her time as student assistant. In addition, we thank various funders for their support: H.H. received funding from the Else Kröner Fresenius Foundation, Bad Homburg, Germany, that covered the costs for conducting the HuMet study including collection and storage of the biosamples. K.S. is supported by the Biomedical Research Program at Weill Cornell Medicine in Qatar, a program funded by the Qatar Foundation. M.A. and G.K. received funding (through their institutions) from the National Institutes of Health/National Institute on Aging through grants RF1AG057452, RF1AG058942, RF1AG059093, U01AG061359, U19AG063744, and R01AG069901. J.R. was funded by grant 01ZX1912D by the BMBF within the framework of e:Med research and funding concept. This work was supported by the de.NBI Cloud within the German Network for Bioinformatics Infrastructure (de.NBI) funded by the German Federal Ministry of Education and Research (BMBF) (031A532B, 031A533A, 031A533B, 031A534A, 031A535A, 031A537A, 031A537B, 031A537C, 031A537D, 031A538A).

## Data availability

The datasets for this study can be found in the Download section of the HuMet Repository: https://humet.org. Data from the non-targeted metabolomics platform is also available at the MetaboLights Database: http://www.ebi.ac.uk/metabolights/MTBLS89.

